# Quantifying oligomer populations in real time during protein aggregation using single-molecule mass photometry

**DOI:** 10.1101/2022.06.15.496189

**Authors:** Simanta Sarani Paul, Aaron Lyons, Russell Kirchner, Michael T. Woodside

**Author notes:** These authors contributed equally (Simanta Sarani Paul, Aaron Lyons).

## Abstract

Protein aggregation is a hallmark of many neurodegenerative diseases. The early stages of the aggregation cascade are crucial because small oligomers are thought to be key neurotoxic species, but they are difficult to study because they feature heterogeneous mixtures of transient states. We show how the populations of different oligomers can be tracked as they evolve over time during aggregation by using single-molecule mass photometry to measure individually the masses of the oligomers present in solution. Applying the approach to tau protein, whose aggregates are linked to diseases including Alzheimer’s and frontotemporal dementia, we found that tau existed in an equilibrium between monomers, dimers, and trimers before aggregation was triggered. Once aggregation commenced, the monomer population dropped continuously, paired first with a rise in the population of the smallest oligomers and then a steep drop as the protein was incorporated into larger oligomers and fibrils. Fitting these populations to kinetic models allowed different models of aggregation to be tested, identifying the most likely mechanism and quantifying the microscopic rates for each step in the mechanism. This work demonstrates a powerful new approach for characterizing previously inaccessible regimes in protein aggregation and building quantitative mechanistic models.

---

A wide range of neurodegenerative diseases, including Alzheimer’s, Parkinson’s, Huntington’s, prion diseases, ALS, and frontotemporal dementia (FTD), feature the progressive accumulation of aggregates of a variety of misfolded proteins as hallmarks of disease.^1-3^ Although these diseases differ in their etiology and presentation, as well as the proteins involved in misfolding, their aggregation cascades show common features: initiated by misfolding events or destabilization of the native structure, the aggregation proceeds through nucleation of a variety of oligomers that grow in size and typically mature eventually into stable amyloid fibrils,^3, 4^ Although large fibrillar aggregates are usually prominent pathological features, in many cases small oligomers formed early in the aggregation cascade are thought to be more important as neurotoxic species.^2^

Extensive efforts have been made to develop therapeutics that act by inhibiting aggregation,^5-7^ but they have been hampered by the technical difficulty of monitoring and studying small oligomers: oligomers typically exist only transiently, as minority sub-populations in a very heterogeneous mixture where most of the protein is usually either natively folded or incorporated into large aggregates like amyloid fibrils. Commonly used methods for monitoring aggregation such as light scattering, thioflavin-T (ThT) fluorescence, or size-exclusion chromatography are generally sensitive to only the largest aggregates (*e*.*g*. fibrils) or else the very smallest (*e*.*g*. dimers/trimers), leaving much of the range of oligomer sizes early in the aggregation cascade undetected. Such methods are often used to infer what happens in the early stages of aggregation based on characterizing the lag time required for nucleation of the large aggregates that are most easily detected.^8^ Measurements of aggregation as a function of protein concentration can be used to build fairly detailed kinetic models of the process.^9^ However, because these approaches do not directly detect oligomers and determine their sizes, the reliability of the models they produce is not always clear.

Methods that monitor aggregation at the single-molecule (SM) level can in principle overcome these challenges for studying aggregation, as they are ideal for characterizing sub-popula-tions of rare or transiently occupied states that would otherwise remain hidden within ensemble averages.^10^ However, the methods used to date all have important technical constraints that limit their ability to quantify oligomer populations. Imaging by electron or atomic force microscopy of proteins deposited on surfaces at different points during the aggregation cascade can give a qualitative sense of how the ensemble of aggregated species changes over time, but precise quantification of oligomer size from the length and/or area in images is often unreliable.^11, 12^ SM force spectroscopy can detect and characterize a wide variety of structures formed in small oligomers of a given size,^13, 14^ but it has limited ability to probe changes in oligomer populations over time. SM fluorescence methods can monitor changes in oligomer sizes and conformations over time using correlation spectroscopies or Förster resonant energy transfer (FRET),^15-17^ but they have difficulty quantifying the distribution of oligomer sizes precisely and require the use of fluorescent dyes that may affect aggregation.

A promising new approach is offered by SM mass photometry (SMMP),^18^ whereby individual molecules are detected from the interference between back-scattered and reflected light as they bind non-specifically to a microscope coverslip (Fig. 1). Because the optical contrast varies linearly with particle volume, it can be used to quantify the mass of individual molecules.^19^ SMMP therefore holds out the promise of determining the distribution of oligomer sizes directly and reliably, label-free. By monitoring changes in the populations of different oligomer sizes over time, the kinetics of each step in the aggregation cascade can in principle be measured. SMMP has been used to explore oligomer formation by bovine serum albumin,^19^ peroxiredoxins,^20^ and MinDE,^21^ as well as the rate of fibril growth in actin.^19^ It has also been used to monitor the growth of α-synuclein fibrils,^19^ but nucleation of oligomers and the individual steps in aggregate assembly could not be detected, preventing a full characterization of aggregation mechanisms.

**Figure 1:**
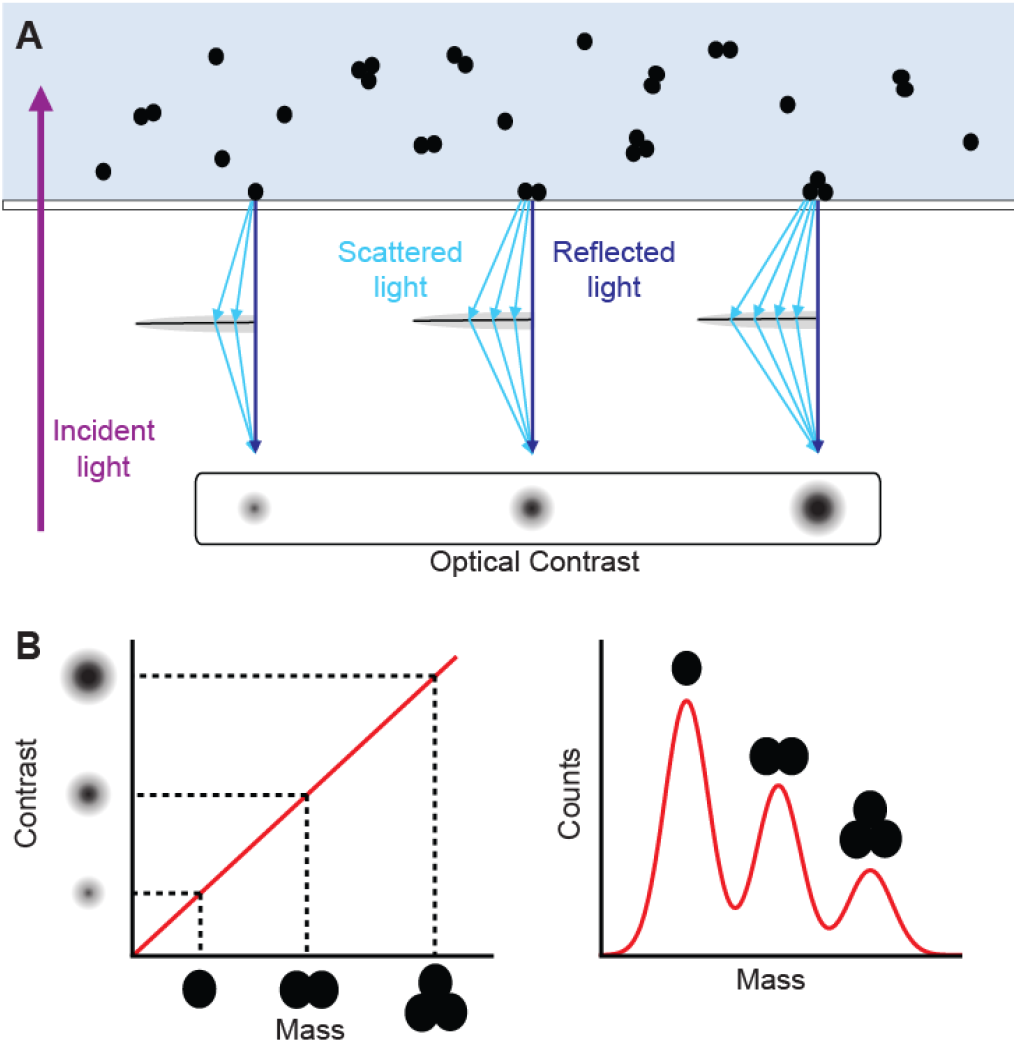
Single-molecule mass photometry. Schematic describing the operating principles of SMMP. (A) Protein molecules binding to a coverslip surface scatter light in a size-dependent way. Interference with the incident light reflected from the coverslip produces optical contrast that varies with the size of the detected protein. (B) The linear relationship between optical contrast and protein mass is used to determine the mass of the object in each binding event (left) and measure mass distributions (right).

Here we demonstrate how SMMP can be used to quantify the populations of oligomers of different size, and thereby test and fit kinetic models of aggregation, by studying the protein tau. Aggregates of misfolded tau are characteristic features in a wide range of neurodegenerative diseases, including Alzheimer’s disease, frontotemporal dementia, chronic traumatic encephalopathy, Pick’s disease, and progressive supranuclear palsy,^22^ making tau aggregation an important diagnostic and therapeutic target. We use SMMP to count the number of oligomers with sizes ranging from monomers to 25-mers. We find that under conditions where tau does not aggregate, it maintains an equilibrium between monomers, dimers, and trimers. Triggering aggregation, however, induces the formation of higher-order oligomers, which first accumulate after an initial lag phase before rapidly disappearing as large-scale aggregates are formed. By modeling the kinetics for oligomers of different size, we solve for the aggregation mechanism and quantify all the microscopic rates, finding that tau aggregates slowly and reversibly until hexamer formation, at which point it undergoes rapid and near-irreversible elongation. This work demonstrates the ability of SMMP to elucidate the mechanisms of protein aggregation by tracking and quantifying individual oligomer populations, providing a detailed account of the kinetics governing the earliest steps of the aggregation cascade.

## RESULTS

### Native tau is in a monomer-dimer-trimer equilibrium

We first examined the oligomeric state of tau under conditions when it does not aggregate, studying specifically the 0N4R isoform of human tau. Wild-type tau is generally resistant to spontaneous aggregation in the absence of agents that induce aggregation, likely due to repulsion between positively charged residues.^23^ We therefore measured it in aqueous buffer without any inducer present. Analyzing the distribution of masses seen during a 1-min recording of the scattering from molecules landing on the coverslip surface, we found three peaks (Fig. 2A): the dominant one at 46.7 ± 0.3 kDa, a secondary peak at 86.2 ± 0.7 kDa, and a minor peak at 129 ± 1 kDa, where the uncertainties represent the error from fitting the average of four replicates. Given the 43-kDa mass of 0N4R monomers, these peaks are consistent with monomers, dimers, and trimers of tau, respectively. Infrequent observations of objects at higher masses (Fig. 2A, inset) reflect impurities in the protein preparation or possibly higher-order tau oligomers. The mass distribution seen by SMMP remained unchanged after 30 hr of incubation at 37 °C with constant shaking (Fig. 2B), showing no evidence of aggregation (as expected in the absence of inducer) and implying that tau exists in a stable equilibrium between monomers, dimers, and trimers. Determining the fractional occupancy of the different species from Gaussian fits to the peaks, we found that the relative populations of 72 ± 2% monomer, 21.2 ± 0.8% dimer, and 6.6 ± 0.4% trimer suggest the dimer is 0.62 ± 0.08 kcal/mol less stable than monomers on their own at room temperature, whereas the trimer is 2.3 ± 0.1 kcal/mol less stable.

**Figure 2:**
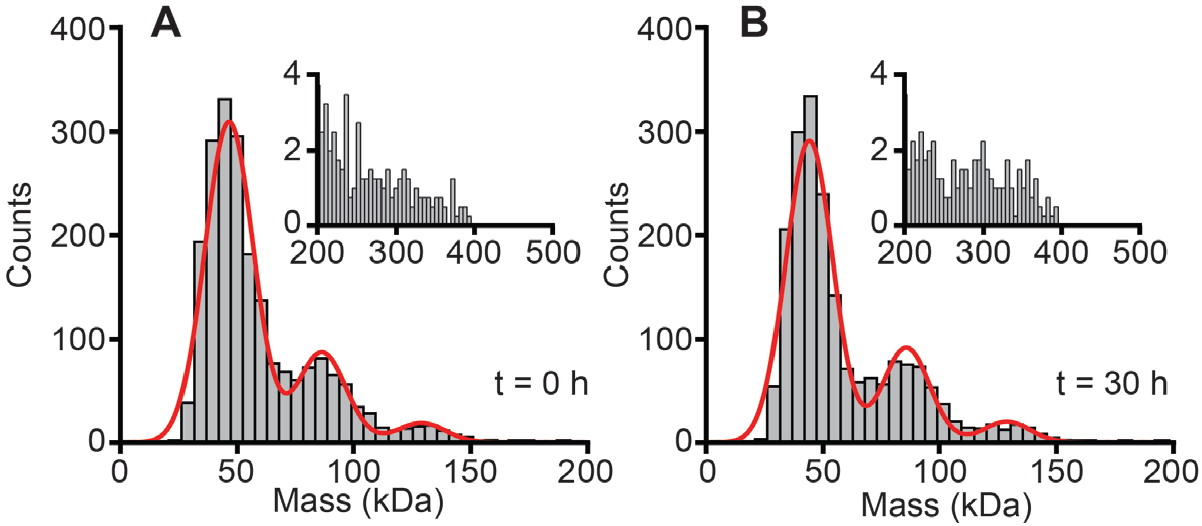
SMMP of tau. (A) On its own, wild-type tau is primarily monomeric, but the mass distribution shows minor peaks corresponding to dimers and trimers. Inset: a small number of binding events at larger mass may represent impurities or rare higher-order tau oligomers. (B) The mass distribution is effectively unchanged after 30-hr incubation at 37 ºC while shaking, indicating wild-type tau does not aggregate spontaneously.

### SMMP monitors oligomer growth during aggregation

Next, we induced aggregation by adding Congo red, a polyanionic azo dye known to induce tau fibrillization,^24^ to the protein solution. Using bulk light scattering to confirm the formation of large-scale aggregates, we saw the characteristic sigmoidal increase in scattering after a lag phase of ∼40 min, reaching saturation after ∼150 min (Fig. 3A); transmission electron microscopy confirmed that aggregation led to the formation of amyloid fibrils (Fig. S1). Turning to the results from mass photometry, unlike in the case without Congo red (Fig. 2), here we saw the oligomer populations change from the initial distribution (Fig. 3B) as aggregation proceeded, with the appearance of higher-oligomers having masses up to the MDa range (Fig. 3C). Distinct shifts in the mass distribution were observed over time (Fig. 3D): the monomer population declined continuously after the first ∼10 min, whereas the dimer and trimer populations increased modestly in the first ∼30 min before declining sharply, and larger oligomers began to appear in the distribution after ∼15 min before declining sharply after ∼40–50 min. The rise and fall in each oligomer population occurred at systematically slightly later times for larger oligomers. Distinct peaks for oligomers up to pentamers and occasionally hexamers could be observed directly in the mass distributions. Increases in the counts at higher molecular weights indicating higher-order oligomers were also detected (Fig. 3C, inset), with examples seen up to ∼25-mers (Fig. 3E), but the counts were insufficient to discern well-defined peaks. Very large aggregates (possibly fibrillar) were also observed towards the end of the measurement, but they could not be imaged reliably to determine their mass from the light scattering. We note that SMMP is relatively insensitive to protein shape, hence the peaks in the mass distribution from native vs misfolded oligomers cannot be distinguished.

**Figure 3:**
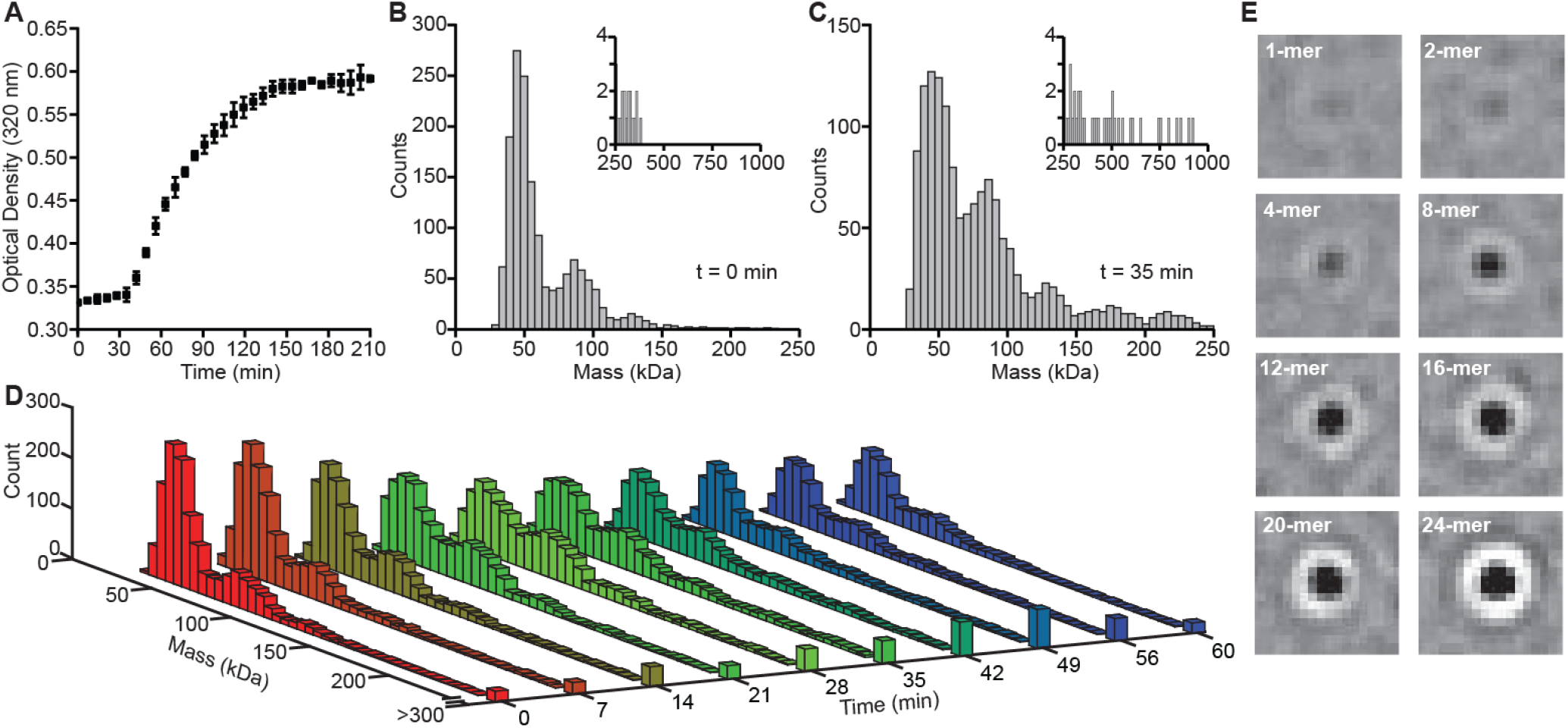
SMMP of tau aggregation. (A) Incubation with Congo red induces aggregation of tau, causing a rapid increase of optical density from light scattering after ∼40 min and leading to the formation of amyloid fibrils. Error bars represent standard error on the mean from 3 replicates. (B) Immediately after addition of Congo red, the mass distribution is the same as seen without inducer. (C) After 35 min of incubation, the prevalence of monomers is decreased while mass peaks corresponding to tetramers and pentamers are seen, as well as binding events at masses up to the MDa range (inset). (D) Mass distributions collected at different times after induction of aggregation show the time evolution of the oligomer populations. (E) Sample optical contrast images of binding events with masses corresponding to the range from monomers and dimers (top) through to 20-mers and 24-mers (bottom). All images are on the same contrast scale, with area 1.43 × 1.43 μm.

To quantify the changes in the oligomer populations as a function of time, we fit the areas of the peaks in the mass distribution corresponding to monomers through pentamers (Fig. 4A) to find the total counts for each species, as illustrated in Fig. 4B. Because the occurrence of oligomers larger than pentamers was too infrequent for reliable fitting of the distributions, we combined the counts from 6-mers to 25-mers into a single population. The number of times each of these oligomeric species was observed was then plotted as a function of the time after aggregation was induced (Fig. 4C) to show how the oligomer population evolved during the aggregation cascade.

**Figure 4:**
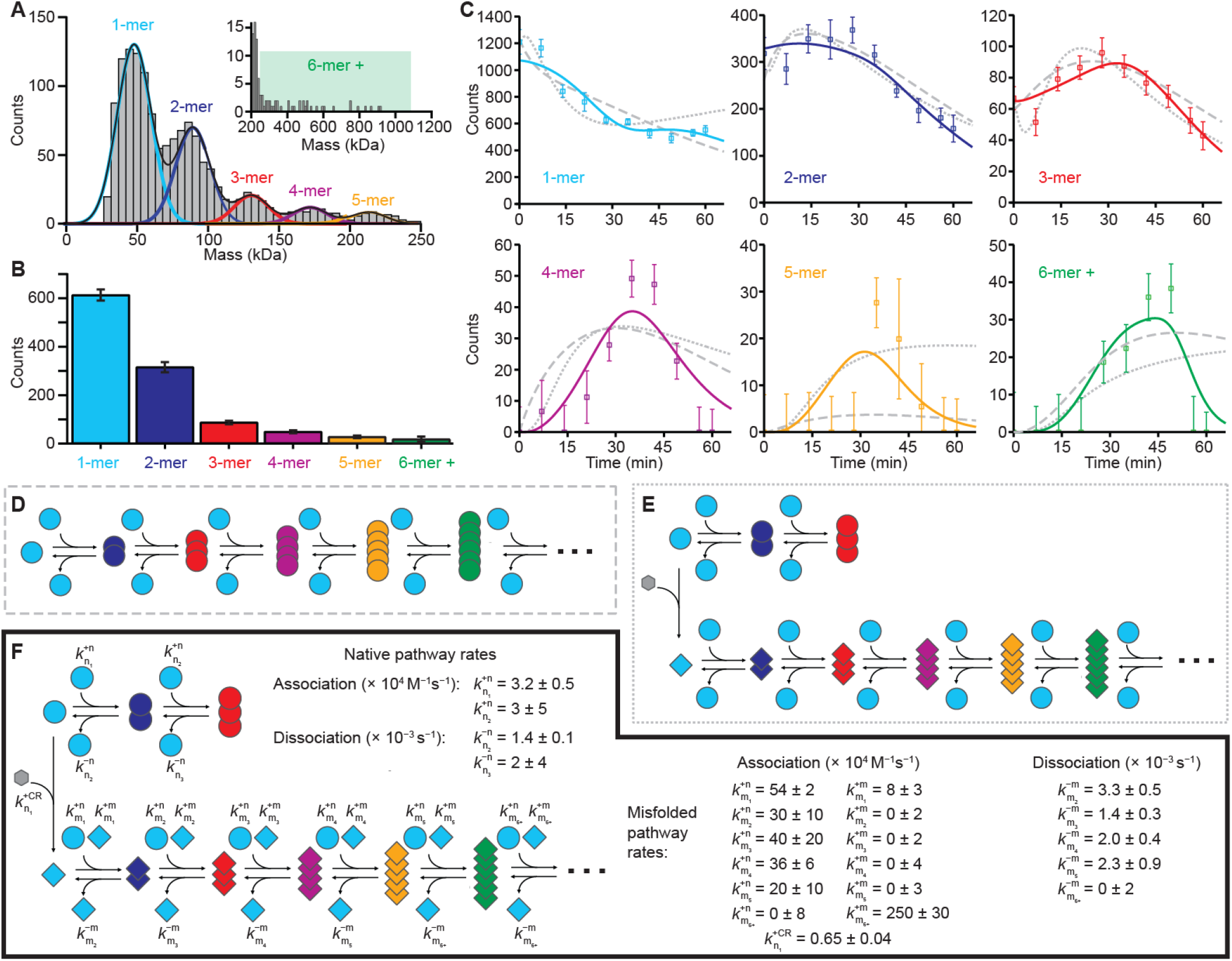
Modeling the aggregation mechanism. (A) Peaks in the mass distribution associated with different oligomers are fitted to determine the number of each oligomer. Inset: because higher-order oligomers are rarer, their counts are integrated across the range of 6-mers to 25-mers. (B) Mass distributions are converted into populations of oligomers. Error bars represent fitting/counting errors. (C) Time evolution of average populations for monomers (cyan), dimers (blue), trimers (red), tetramers (purple), pentamers (orange), and hexamers or larger (green). Error bars represent standard error of mean from three replicates. Dashed lines: fits to model in panel D. Dotted lines: fits to model in panel E. Solid lines: fits to model in panel F. (D) A model assuming a single pathway for aggregation fails to capture the lag phase in the growth of higher-order oligomers. (E) Including an aggregation-incompetent pathway corresponding to the native oligomers observed in the absence of an inducer (circles) as well as an aggregation-prone pathway initiated by the association of native monomers with Congo red (grey hexagon) to form misfolded conformers (diamonds) captures the lag phase but is unable to replicate the rapid drop in large oligomers after ∼45 min. (F) A model where monomers can add to growing oligomers in either native or misfolded conformers but can only disassociate from misfolded oligomers in misfolded conformers captures all of the features of the time evolution data. The monomer addition rate increases dramatically after the formation of pentamers, concomitant with the growth becoming near-irreversible.

### Kinetic modeling reveals microscopic rates for each step in cascade

These direct measurements of the changes in the concentrations of oligomers of specific size allowed us to build and test microscopic kinetic models of the aggregation, thereby measuring the microscopic rates. Such models can give critical insight into aggregation mechanisms, but it is difficult to obtain reliable microscopic rates for each step of the process without quantifying the populations of each oligomeric species. The wealth of kinetic information present in the curves in Fig. 4B—reflecting the growth and decay for each different oligomer size—allow detailed kinetic models of the aggregation to be built and tested, to determine the minimal mechanism that best accounts for the observed aggregation dynamics. We first tested the simplest possible model, which assumes that oligomers lie on a single aggregation pathway and that tau does not change conformation (Fig. 4D). This model was inconsistent with the fact that the dimers and trimers formed by native tau did not lead to higher-order oligomers in the absence of aggregation inducer; not surprisingly, it did not adequately capture the lag time in growth of the oligomers, nor their rapid drop-off with time (Fig. 4C, dashed lines). We next tested a model with two oligomerization pathways, one for native oligomers and the other for non-native oligomers, with the latter initiated by formation of a Congo red-bound non-native monomer; here we assumed that oligomers change size via addition/removal of native monomers (Fig. 4E). This model also did not fully capture the observed dynamics, missing the lag phase in dimers and the rapid drop-off of oligomers larger than trimers after ∼40–50 min (Fig. 4C, dotted lines).

To obtain a model that includes a lag phase for dimer formation, as observed experimentally, we modified the kinetic scheme to assume that any monomers dissociating from misfolded oligomers remain non-native, and to allow for growth by addition of either native or non-native monomers (Fig. 4F). We verified experimentally that Congo red will indeed stay bound to tau monomers, as required for this model (Fig. S2). With the addition of these features to the mechanism, the model captured all the essential features of the oligomer population kinetics (Fig. 4C, solid lines), fitting the data considerably better even when accounting for the increased number of fitting parameters as assessed by the Akaike information criterion (AIC). A key feature of the fit to the kinetics is that the growth of smaller oligomer (dimers to pentamers) is dominated by the slow addition of native monomers, whereas the growth rate of larger oligomers (hexamers and up) is dominated by the much faster addition of misfolded monomers, leading to run-away growth of ever-larger oligomers that rapidly depletes the population of smaller oligomers. Notably, the timing of this rapid growth in larger oligomers, starting ∼40 mins into the aggregation reaction, also coincided with the end of the lag phase in the bulk scattering experiments and the concomitant rapid growth in scattering (Fig. 3A). Interestingly, although oligomers larger than 25-mers were not included in the analysis because they could not be counted reliably, the amount of tau incorporated in such large aggregates could be tracked implicitly through mass conservation by assuming that any missing tau formed large aggregates. We found good agreement between the missing mass and the mass of large aggregates predicted by the model fit (Fig. S3), further supporting the mechanism in Fig. 4F.

We note that including extra steps in the model such as oligomer growth through dimer addition (as well as monomer addition) did not significantly improve the fit and was thus unnecessary to account for the observed kinetics. Attempting to simplify the model by assuming that the on and off rates were independent of the size of the oligomer led to poor fits that were unable to account for the rapid drop-off in oligomers after 40–50 mins (Fig. S4), indicating that size-dependent rates are an essential feature of the kinetic model. However, the results from the model in Fig. 4F indicate similar rates for misfolded dimers through pentamers, suggesting that the model could be simplified by assuming identical rates for this subset of oligomers. This simplification fits only slightly less well than the full model (Fig. S5) but is judged more likely by AIC as the most parsimonious explanation of the data; it is only through the results of the full size-dependent model in Fig. 4F, however, that this minimal model can be identified.

## DISCUSSION

The great advantage of SMMP over other methods for monitoring aggregation is that SMMP allows the population of each different oligomer size to be quantified directly, opening up a new window on the microscopic mechanisms of protein aggregation. The single-molecule nature of the assay is central to this ability, as it enables reliable mass distribution measurements unbiased by differentially size-sensitive detection or labels that may alter the behavior of the protein. Such information is essential for building mechanistic models of aggregation focused on the small oligomers that are likely key neurotoxic actors: the complexity of aggregation reactions, with their potential for multiple pathways and a profusion of different microscopic rates, makes it difficult to build reliable models using assays are not directly and/or uniformly sensitive to all the oligomeric species present. For example, measurements using dyes like thioflavin T are sensitive only to some types of aggregates, measurements using scattering or diffusion time are dominated by the responses from the largest aggregates in the mixture (again leading to biases in oligomer detection), and SM fluorescence approaches have difficulty sizing oligomers owing to the effects of non-uniform excitation in the volume being monitored. Furthermore, because SMMP measures a growth/decay curve for each oligomer of different size, it allows the microscopic on/off rates for each oligomer to be found without overfitting the kinetic model.

Considering the example of tau aggregation studied here, direct counting of oligomers reveals two distinct pathways: one involving oligomerization of native-like protein, which does not lead to larger-scale aggregates and fibrils, and another induced by the aggregation inducer, which does. The aggregation cascade starts with the conversion of monomers into a misfolded, aggregation-competent conformer by Congo red binding, accounting for the observed lag phase in the smallest oligomers. The growth of misfolded oligomers from dimers to hexamers is then relatively slow, involving reversible addition and removal of monomers. However, once the oligomers reach the size of hexamers, their growth becomes effectively irreversible, with the off-rates dropping close to 0 and the on-rates speed up ∼10-fold. This change in the growth rate leads to the accumulation of large-scale aggregates seen in ensemble scattering measurements and accounts for the rapid decrease in all smaller oligomer populations detectable by SMMP. Interestingly, this switch in growth rates coincides with a switch in the source of the growth: the fit to the kinetics finds that the oligomers smaller than hexamers grow almost exclusively by addition of native-like monomers, whereas hexamers and larger oligomers grow by addition of misfolded monomers. The faster growth rate of the larger oligomers may reflect the fact that they no longer need to convert native monomers as in the smaller oligomers. Notably, SMMP shows no evidence of any noticeable accumulation of tau in off-pathway oligomers (at least smaller than 25-mers) other than the native-like dimers and trimers seen in the absence of the aggregation inducer. Curiously, the kinetic modeling suggests that the number of monomers stops dropping briefly around 40–50 mins. Mechanistically, this effect reflects the accumulation of misfolded monomers that are shed from small oligomers, which briefly counteracts the ongoing loss of monomers to form small oligomers until the quasi-irreversible growth of larger oligomers takes off and causes the number of monomers to drop once again.

These results are instructive in helping to clarify the sometimes inconsistent mechanisms proposed for tau aggregation based on previous studies using different methods. Some work has suggested that off-pathway oligomerization is necessary to describe the global kinetics of tau aggregation.^25, 26^ Specifically, direct observation of tau oligomers using FRET demonstrated the presence of two kinetically distinct populations, one that underwent rapid association/disassociation and another with slower kinetics that was hypothesized to be off-pathway.^26^ These findings are similar to our results, suggesting the slow, off-pathway population is comprised of the native-like oligomers observed even in the absence of an aggregation inducer. An additional point of contention in previous studies is the apparent critical nucleus size for starting the aggregation cascade, *i*.*e*. the minimum oligomer size not in equilibrium with the monomer.^27^ Although this nucleus is often assumed to be a dimer^28, 29^, estimates range from monomers^30^ to tetramers.^31^ The challenge with determining the critical nucleus size from experiments like those using ThT fluorescence is that any oligomer of the critical nucleus size or higher is assumed to be fibrillar and thus observable: this assumption couples the results of the kinetic modelling to resolution limit of the experiment and may explain some of the variability seen in critical nucleus size estimates. In contrast, our results—which are uniquely sensitive to small oligomers—suggest that the smallest oligomer not in equilibrium with the monomer is the hexamer. We also find that the rate-limiting step for aggregate growth is not the addition of monomers to the oligomer preceding this critical nucleus, an assumption that is required for certain functional forms describing fibril growth. Rather, the rates of monomer addition and removal are comparable for the dimer through to the pentamer (Fig. 4F), suggesting that ascribing a single rate-limiting step may not be appropriate in the case of tau aggregation. It is difficult to compare the rate constants from our analysis directly to those in the literature, because of differences in the measurement conditions and kinetic modeling, but the on rates found here are similar in magnitude to those reported^11^ for tau aggregation induced by Thiazine red (∼10^5^ M^−1^s^−1^) whereas the off rates are somewhat slower (∼10^−2^ M^−1^s^−1^, from disaggregation measurements).

As a powerful tool for distinguishing small oligomers, SMMP has numerous potential applications in studying aggregation. In addition to deciphering key features of the microscopic mechanisms of neurodegeneration-related aggregation, as illustrated above, SMMP should also be able to probe how pathogenic mutations or post-translational modifications^32-34^ alter these mechanisms, providing new insight into the parts of the cascade that may be most dangerous for disease. Furthermore, SMMP should prove a useful tool for testing potentially therapeutic aggregation inhibitors, not only detecting directly if they inhibit the formation of toxic small oligomers but also determining their mechanism of action by quantifying the effects on each step in the cascade. SMMP does face some technical challenges for studying aggregation. For example, being sensitive to all objects that scatter light, it requires high-purity samples. This consideration also limits what can be used to induce aggregation: for example, heparin is commonly used as an inducer for tau,^35^ but its large mass broadens the observed mass distributions, obscuring the peaks from different oligomers. A preponderance of small oligomers can also make it difficult to obtain sufficient counts to define the populations of larger oligomers and fibrils reliably, although this difficulty can be mitigated by amalgamating the counts from a range of oligomer sizes (as done here) or by increasing the number of measurements. Combining SMMP with other methods may offer the most promising approach: by first establishing the mechanism of early aggregation, SMMP data could provide unique insight complementing the results from traditional assays of fibril growth. Such a combined approach might also clarify mechanistic features of late-stage aggregation processes like secondary nucleation and fragmentation^3^, yielding a more global picture of protein aggregation.

## METHODS

### Sample preparation

The 0N4R isoform of tau was purified as described previously^36^. Briefly, 0N4R tau was expressed in BL21-RP cells transfected with the vector pMCSG7, inducing expression by adding IPTG at 300 μL/L culture at 30 °C for 2 hr. Cell pellets formed in high-speed centrifugation were resuspended at 15 mL/L in 20 mM 2-(N-morpholino)ethanesulfonic acid (MES) buffer, pH 6.8, containing 1 mM ethylene glycol-bis(β-aminoethyl ether)-N,N,N′,N′-tetraacetic acid (EGTA), 0.2 mM MgCl_2_, 1 mM phenylmethylsulfonyl fluoride (PMSF) protease inhibitor, and 5 mM dithiothreitol (DTT) at 4 °C. Cells were lysed with a microfluidizer at 13,000 kPa under ice-cold conditions, then the diluted lysate was boiled for 20 minutes to denature all proteins except tau. The cell debris, other denatured proteins, and salts were removed by centrifugation and dialysis in 20 mM MES buffer, pH 6.8, containing 50 mM NaCl, 1 mM EGTA, 1 mM MgCl_2_, 0.1 mM PMSF and 2 mm DTT. Finally, tau was purified by cation-exchange chromatography (SP sepharose in XK26 column). Tau concentration was measured by BCA assay.

### SMMP measurements

SMMP measurements were done with the OneMP mass photometer (Refeyn), calibrated as per manufacturer protocols using bovine serum albumen (BSA) and apoferritin. Calibration error was estimated at 3%. All measurements involved a total volume of 15 μL in the measuring well, first adding 10–13 μL of measuring buffer to clean wells before adding protein to a final concentration of 40 nM. To ensure cleanliness, buffers for SMMP measurements were filtered twice with 0.22-µm syringe filters; buffer cleanliness was also assayed before each measurement run by taking a 1-min SMMP measurement of buffer only to verify that the buffer did not contain any particles generating optical contrast.

For measurements of tau under non-aggregating conditions, tau protein was suspended at a final concentration of 10 μM in phosphate buffered saline (PBS), pH 7.5, with 1 mM PMSF and 2 mM DTT, and incubated at 37 °C while shaking at 300 rpm for 24–30 hr. Samples collected at 0 and 24 or 30 hr of incubation were serially diluted to 120 nM in PBS at room temperature before being diluted to the final concentration of 40 nM in the sample well. Movies of the scattering were recorded for 1 min and analyzed as described below. For measurements of aggregation, tau protein was suspended at a final concentration of 10 μM in phosphate buffered saline, pH 7.5, with 1 mM PMSF and 2 mM DTT, and incubated at 37 °C while shaking at 300 rpm, as above. Aggregation was then initiated in the solution, prior to any measurements using SMMP, by adding Congo red at a final concentration of 5 μM. At designated time-points, samples were collected from the stock aggregation solution, then diluted to 500 nM and placed on ice to slow down the aggregation reaction, before being serially diluted 40 nM at room temperature in the sample well. For each time-point in the aggregation, movies of the scattering were recorded for 1 min and analyzed as described below. Three biological replicates were done of all measurements.

### SMMP data analysis

Movies of the scattering was analyzed using DiscoverMP software (Refeyn), using as default 10 binned frames and a median filter kernel of value 1.5. Ratiometric images yielding the change in the light scattering between image frames were analyzed to find binding events using default thresholds for intensity change and sensitivity. The image from each binding event was fitted to a point spread function to find the center of the particle. The optical contrast value at the particle center was then compared to the calibration curve obtained from measurements of BSA (monomers: 66 kDa; dimers: 132 kDa) and apoferritin (480 kDa) to determine the mass of the particle. The distribution of all particle masses from each 1-min run was then plotted. Note that the only aggregates detected by SMMP are those that are already formed in solution before the measurement; aggregates that nucleate on the glass surface are not seen directly, since the ratiometric analysis limits the detection to mass that is added to the surface (*i*.*e*., only monomers in solution that are incorporated into a putative surface-bound aggregates are seen, not the surface-bound aggregate itself).

For measurements without Congo red, histograms of the mass distribution (5.2-kDa bin size) were fit between 26 and 197.6 kDa to the sum of three Gaussian distributions. The standard deviations were constrained to be equal, but the relative peak positions were unconstrained to test the hypothesis that the peaks were located at integer multiples of the tau mass. The free energy differences for the native monomer, dimer, and trimer were found by calculating the fraction of tau monomers that occupied each of these states (based on the counts for each state multiplied by the number of monomers in each state) and applying the Boltzmann formula to determine the energy difference per monomer. The dimer stability was then twice that of each monomer in the dimer, and the trimer stability thrice that of each monomer in the trimer.

For measurements where aggregation was induced by Congo red, histograms from measurements at a given time after addition of Congo red were fit between 26 and 239.2 kDa to the sum of five Gaussian distributions, corresponding to monomers through pentamers, which were constrained to have equal standard deviations and inter-peak spacing. The area under each Gaussian was then calculated to determine the total counts of each of these oligomers. For the population of 6-mers to 25-mers, the total counts observed between 241.8 and 1094.6 kDa were integrated. Because the low background of large-mass events seen in the absence of Congo red suggests a background signal from sample impurities, we subtracted this background from the total counts for each oligomer, estimating the background from a linear fit to the averaged bin counts at 200–400 kDa in the absence of Congo red. This calculation was repeated for each of the three replicates, and the total counts for each oligomer species averaged at each time-point. All fitting was done with Igor Pro 8.0.4.

Kinetic models were coded as systems of first-order differential equations and solved using the ode45 function in MATLAB R2021a (MathWorks, 2021). Equations for each model are listed in Supplementary Information. After a coarse search for starting points for the kinetic parameters, optimal parameters were found using the Levenberg-Marquardt algorithm (lsqnonlin function in MATLAB) to minimize the chi-squared error between the model and data. Once the algorithm had converged, we used the Jacobian matrix at convergence to estimate the uncertainties on the optimal parameters^37^: *σ*_*i*_ = ([*J*^T^*WJ*]_*ii*_)^1/2^, where *σ*_*i*_ is the estimated standard deviation of the *i*th fit parameter at convergence, *J* is the *n* × *m* Jacobian matrix at convergence (with *n* the number of data points being fit and the number of fit parameters in the model), and *W* is the diagonal weight matrix whose entries are the inverse variances of the data points used in the fit. For the models that explicitly included Congo red, the initial number of Congo red molecules was set at half the total number of tau monomers at zero time, consistent with the experimental conditions. We converted count units from the experiment/modeling into molar units by determining the total number of tau monomers that bound to the surface over the course of the measurements without Congo red and using the linear relation between protein concentration and counts, finding a conversion factor of 83 ± 7 counts/nM. The relative likelihood that each of the 3 kinetic models outlined in Fig. 4 explained the observed data was assessed with the Akaike information criterion, AIC. We found ΔAIC < −120 for the third model compared to the first and second models, indicating that the third model is orders of magnitude more likely than the other two. A fourth model that expanded on the model in Fig. 4F by including the possibility of oligomer growth via addition of misfolded dimers was also tested, but it had ΔAIC = +9.6 compared to the model from Fig. 4F, indicating that it was only 0.8% as likely as the latter. For the model that assumes size-independent on/off rates (Fig. S4), ΔAIC = +126.5 compared to the model from Fig. 4F, indicating that it is extremely unlikely, whereas ΔAIC = −14.6 for the model assuming the same on/off rates for dimers through pentamers (Fig. S5), suggesting that it is the minimal model accounting for the data.

### Ensemble scattering measurements

Samples were collected at specific time-points during the aggregation process and the scattering measured in a spectrophotometer (Agilent Cary 60) at 320 nm.

## Acknowledgements

We thank Allan Yarahmady and Sue-Ann Mok for generously sharing the plasmid used to express tau, as well as for sharing expertise and equipment for purifying tau. This work was funded by Alberta Innovates (grant number G2018000860) and the Natural Sciences and Engineering Research Council of Canada (grant number RTI-2020-00301). AL was supported by an Alberta Innovates Graduate Fellowship.

## Author contributions

SSP and RK prepared samples; SSP and RK collected data; AL developed analysis methods; AL and SSP analyzed data; SSP, AL, and MTW wrote manuscript.

## Supplemental Information

### Equations for kinetic modelling

The differential equations describing the three kinetic models tested in Fig. 4 are listed below.

#### Model 1 (Fig. 4D): single pathway

Here, *n*_*i*_ is the number of oligomers of size *i*, and the various rates *k* describe addition of a native monomer (superscript +n) or removal of one (superscript −n) onto/from an oligomer of size *i* (subscript n_*i*_). Oligomers of size *N* = 50 are taken to be a sink term (*i*.*e*., monomers can be added to them but never removed).

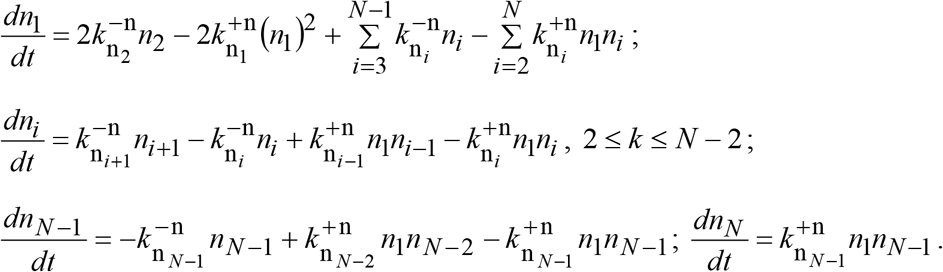

#### Model 2 (Fig. 4E): double pathway, native addition/removal

Here, interaction of native monomers with Congo red produces misfolded monomers, which can grow into misfolded oligomers. Native oligomers (with number *n*_*i*_ as above) exist in a monomer-dimer-trimer equilibrium independent of the misfolded oligomers (with number *m*_*i*_). The rates (*k*) describe addition of a native monomer (superscript +n), addition of a misfolded monomer (superscript +m), or removal of a native monomer (superscript −n) onto or from a native oligomer of size *i* (subscript n_*i*_) or a misfolded oligomer of size *i* (subscript m_*i*_). *k*^+CR^ describes the rate of interaction of monomers with Congo red.

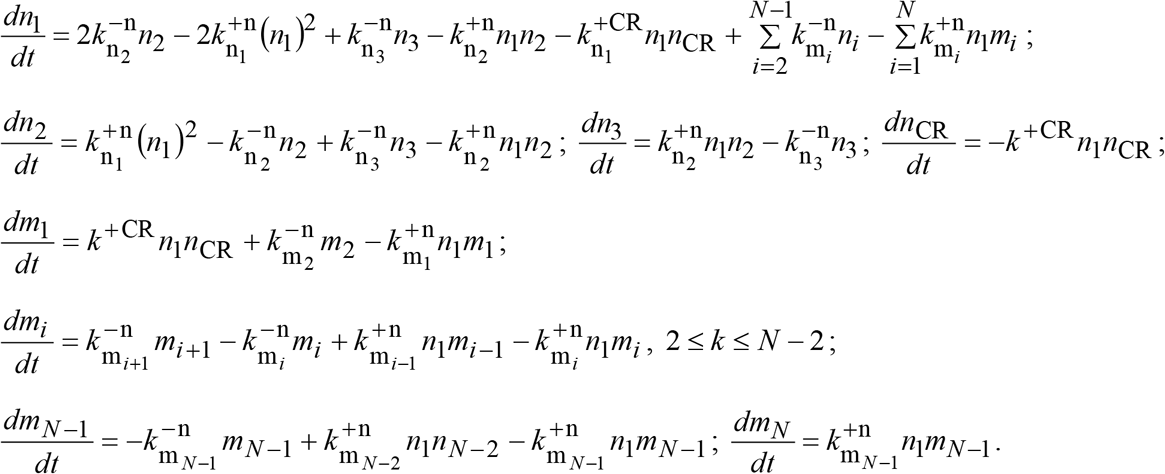

#### Model 3 (Fig. 4F): double pathway, misfolded addition/removal

Here, monomers can add to growing oligomers in either native or misfolded conformers, but when they dissociate from misfolded oligomers they remain misfolded.

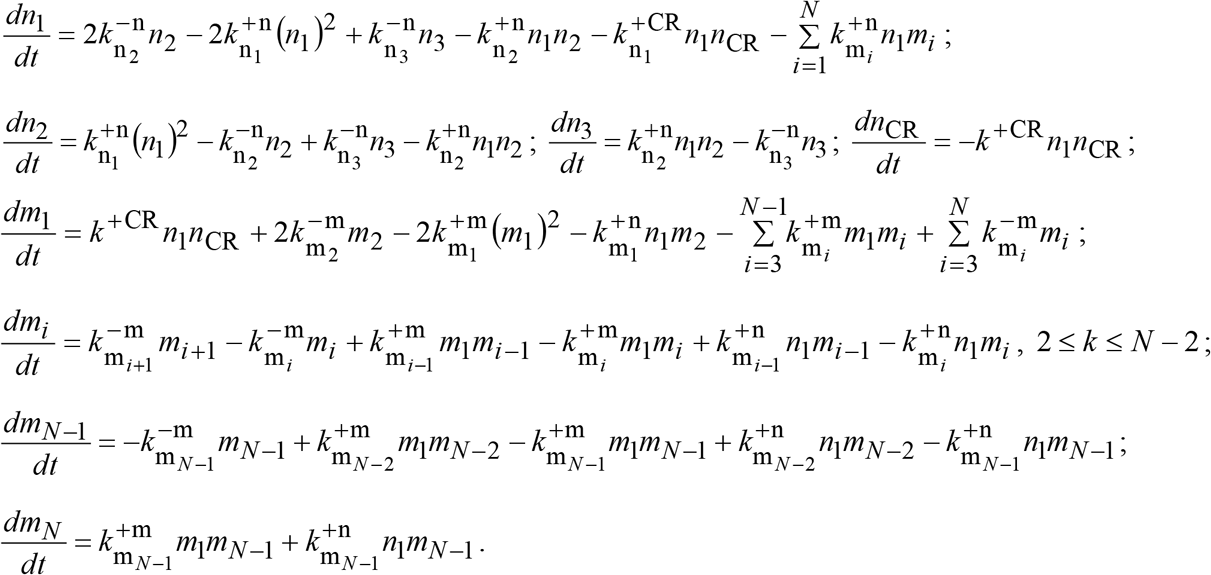

**Figure S1:**
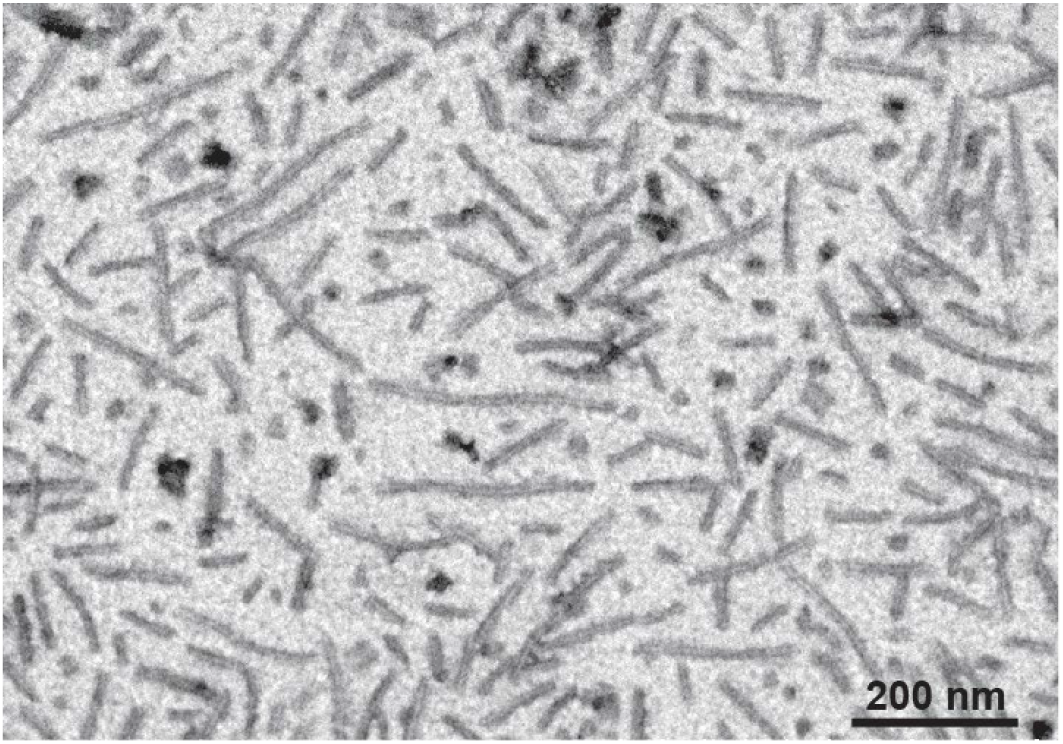
TEM image of amyloid fibrils induced by Congo red. Scanning transmission electron microscope image (Hitachi S-5500) using cold field emission of tau protein incubated with Congo red for 24 hr at 37 °C and 300 rpm shaking (prepared as for SMMP measurements). 5 μL of tau-Congo red sample was deposited onto a TEM grid for 30 s, dried, and then gently washed with 10 µL of deionized water to remove the salt from the aggregation mixture. The grid was stained with 5 µL of 2% aqueous uranyl acetate for 30 s, dried, and imaged at 30 kV accelerating voltage and 30 µA emission current.

**Figure S2:**
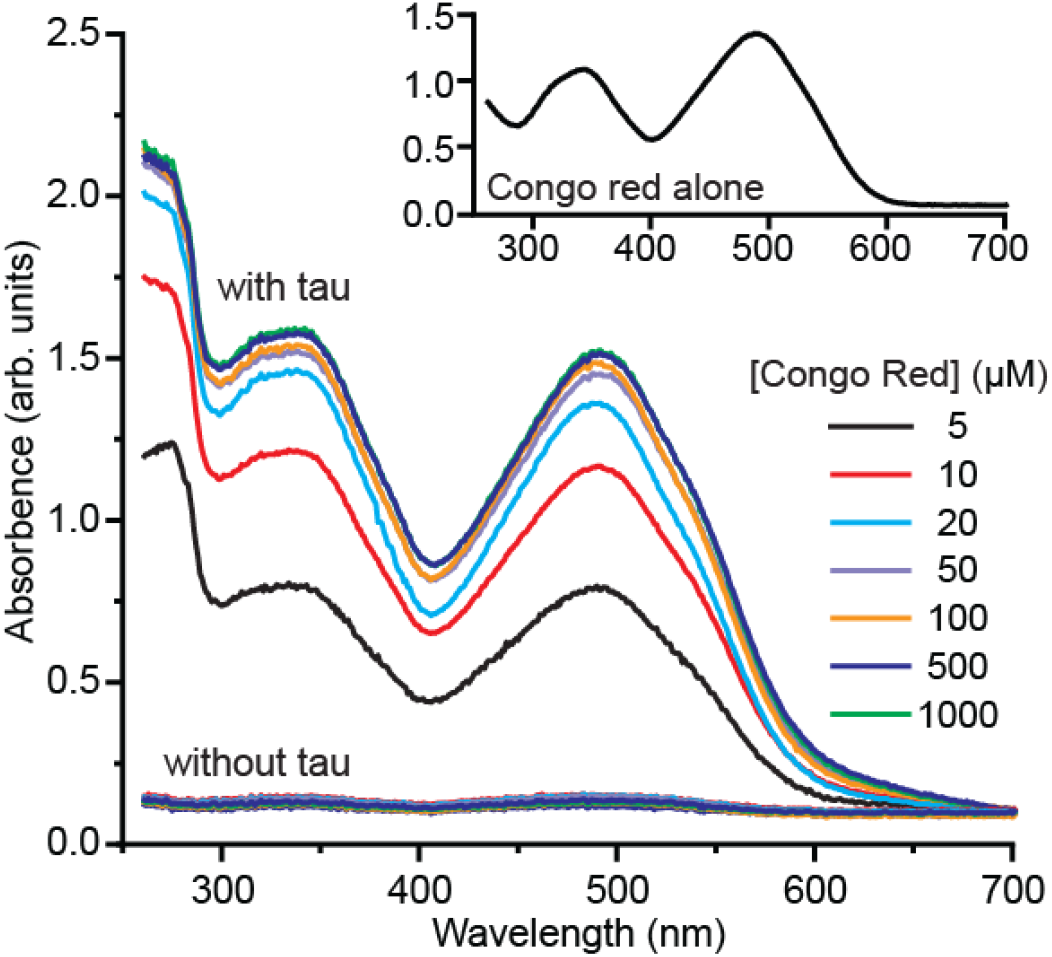
Congo red stays bound to monomeric tau. To verify that Congo red stays bound to monomeric tau, we incubated tau (without Congo red) in 6 M guanidine hydrochloride at room temperature for 2 hr to dissociate native oligomers, immobilized the resulting monomers on a Ni-NTA agarose column, and removed the denaturant and any unbound tau by washing with PBS buffer containing 20 mM imidazole. Equal volumes of column-bound tau and Congo red at various concentrations were mixed and incubated for 30 min at room temperature to allow binding, then the Congo red solution was washed off using 25 column volumes of PBS buffer with 20 mM imidazole. The immobilized tau was then eluted from the column with ice-cold PBS containing 1M imidazole, and the absorbance of the eluted mixture was measured immediately. As a control, the measurements were repeated but without any tau protein present, to measure the background absorbance from any Congo red that bound non-specifically to the column. Inset: Absorbance spectrum of 45 nM Congo red.

**Figure S3:**
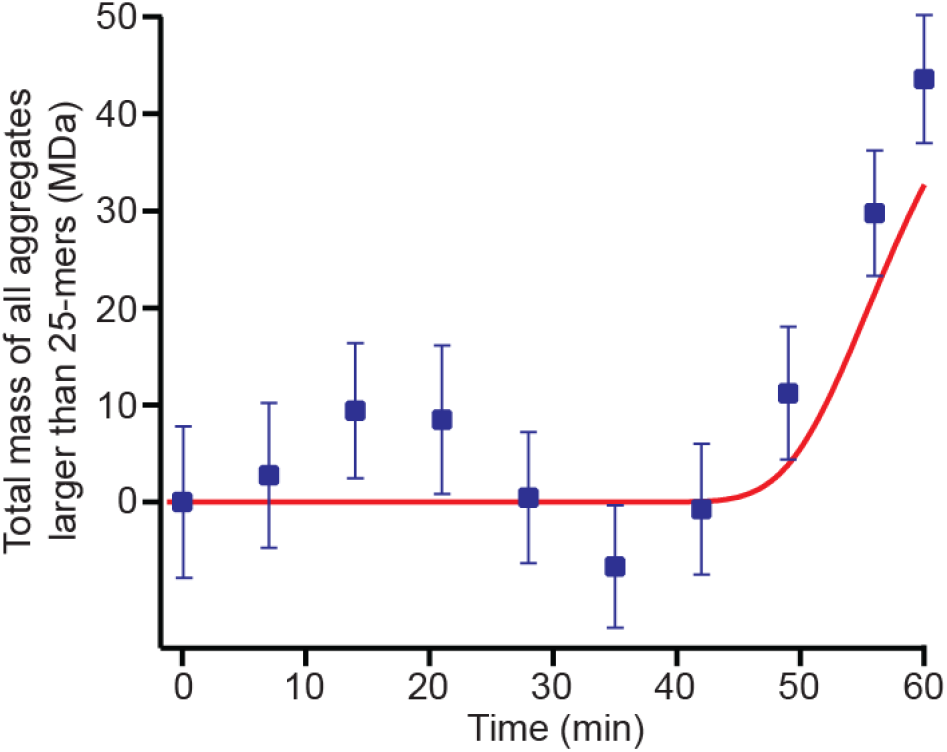
Kinetics of large aggregates inferred from missing mass. The amount of tau mass that appeared to be missing based on oligomer counts (blue), as obtained from the total mass of tau monomers counted per unit time at a given aggregation time-point compared to the total mass of tau monomers seen at 0 s of aggregation, agreed well with the amount of mass expected from the fit to the model in Fig. 4F to be incorporated in aggregates larger than 25-mers (red).

**Figure S4:**
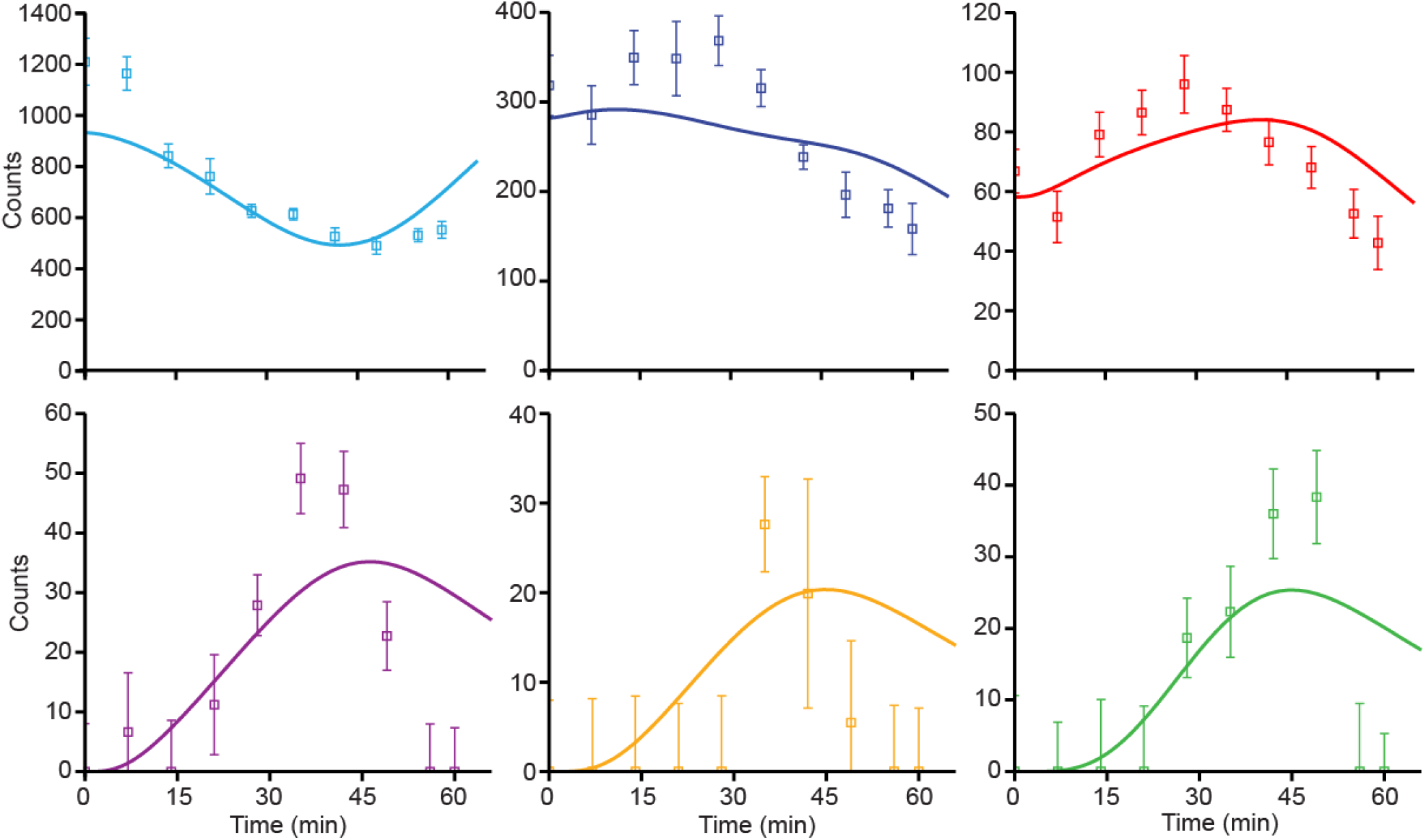
Size-dependent on/off rates are required to fit the oligomer kinetics. Simplifying the model in Fig. 4F by assuming that on and off rates are independent of the oligomer size fails to fit the observed oligomer kinetics.

**Figure S5:**
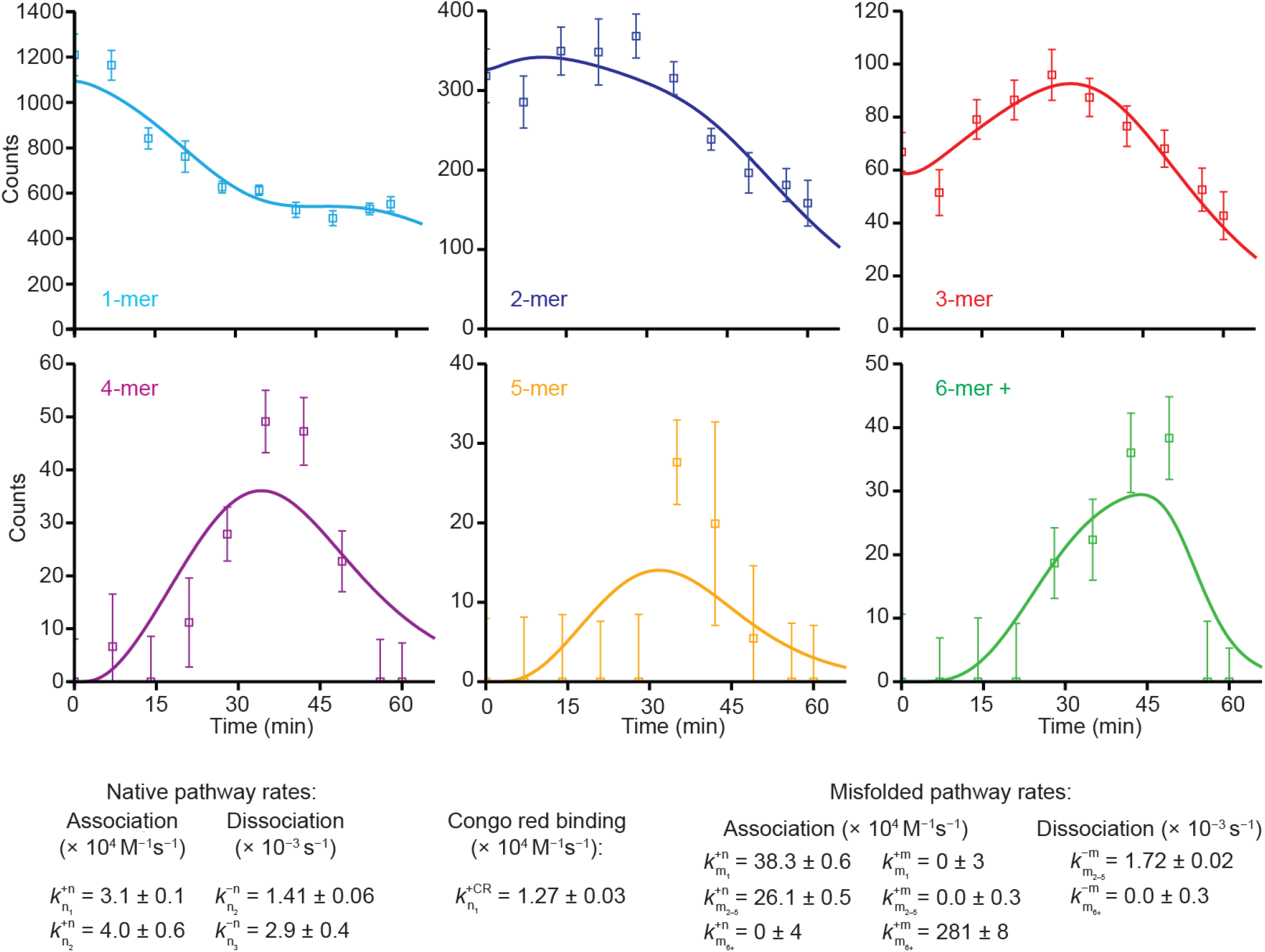
Fit of oligomer kinetics assuming size-independent rates for dimers through pentamers. Simplifying the model in Fig. 4F by assuming that the on and off rates are independent of oligomer size for dimers, trimers, tetramers, and pentamers fits the observed kinetics well.

